# CRISPR/Cas9-mediated gene knockin in the hydroid *Hydractinia symbiolongicarpus*

**DOI:** 10.1101/342592

**Authors:** Steven M. Sanders, Zhiwei Ma, Julia M. Hughes, Brooke M. Riscoe, Gregory A. Gibson, Alan M. Watson, Hakima Flici, Uri Frank, Christine E. Schnitzler, Andreas D. Baxevanis, Matthew L. Nicotra

## Abstract

**Background:** *Hydractinia symbiolongicarpus*, a colonial cnidarian, is a tractable model system for many cnidarian-specific and general biological questions. Until recently, tests of gene function in *Hydractinia* have relied on laborious forward genetic approaches, randomly integrated transgenes, or transient knockdown of mRNAs.

**Results:** Here, we report the use of CRISPR/Cas9 genome editing to generate targeted genomic insertions in *H. symbiolonigcarpus.* We used CRISPR/Cas9 to promote homologous recombination of two fluorescent reporters, eGFP and tdTomato, into the *Eukaryotic elongation factor 1 alpha (Eef1a*) locus. We demonstrate that the transgenes are expressed ubiquitously and are stable over two generations of breeding. We further demonstrate that CRISPR/Cas9 genome editing can be used to mark endogenous proteins with FLAG or StrepII-FLAG affinity tags to enable *in vivo* and *ex vivo* protein studies.

**Conclusions:** This is the first account of CRISPR/Cas9 mediated knockins in *Hydractinia* and the first example of the germline transmission of a CRISPR/Cas9 inserted transgene in a cnidarian. The ability to precisely insert exogenous DNA into the *Hydractinia* genome will enable sophisticated genetic studies and further development of functional genomics tools in this understudied cnidarian model.

## Background

Cnidarians are a phylum of morphologically simple, primarily marine invertebrates. They have long served as experimental models in the fields of developmental biology, symbiosis, and self/non-self recognition (reviewed in [1–4]). More recently, cnidarians have generated interest as models for regenerative and stem cell biology [5–7], neurobiology [8,9], biophysics [10,11] and aging [12,13]. As the closest outgroup to bilaterians [14,15], cnidarians also provide context for our understanding of how fundamental biological processes have evolved in other animals, including humans. Indeed, cnidarian genomes are remarkably similar to the human genome in terms of gene content and structure [16,17]. Cnidarian genomes also possess many human disease genes not found in other animal models [18]. For these reasons, there is an increasing demand for tools allowing direct tests of gene function in cnidarian species.

*Hydractinia* is a genus of colonial marine hydroids and a rapidly maturing model system for cnidarian genetics and functional genomics. Like the two most commonly used cnidarian models, *Hydra* and *Nematostella*, *Hydractinia* has played a key role in our understanding of the molecular basis of development and tissue regeneration [19–22]. Because *Hydractinia* diverged from *Hydra* 306-428 million years ago and *Nematostella* 570-635 million years ago [23–26] (divergence times roughly equivalent to the human/frog and human/hagfish splits, respectively [27]), genetic studies that include these three taxa increase the phylogenetic resolution with which we can understand the evolution of gene function. In addition, *Hydractinia* exhibits traits that are difficult or impossible to study at the molecular level in other cnidarians, including a pluripotent stem cell lineage called interstitial cells (i-cells) [6], coloniality [28], morphological polymorphism [20,29], and allorecognition [4].

From a practical standpoint, *Hydractinia* offers several advantages. It is small in size and inexpensive to culture. Robust laboratory strains for one *Hydractinia* species *(H. symbiolongicarpus)* are available (Nicotra ML, personal communication), and a project to sequence its genome is underway [30]. It is transparent, facilitating *in vivo* microscopic observations throughout the animal at all developmental stages. It is also simple to breed because it spawns daily in response to a light cue, generates embryos that are easy to manipulate, grows into adults that are sessile and dioecious, and has a three-month generation time (comparable to mice and zebrafish). In addition, colonies can be clonally replicated and does not appear to age in the lab, allowing researchers to work with the same animal for years. These features are one reason that *H. symbiolongicarpus* is the only cnidarian in which a forward genetic approach has been used to identify the genetic basis of a phenotype [31,32]. Reverse genetic techniques are also possible in *Hydractinia*, including RNA interference (RNAi) [21,33,34], morpholinos [35], and the creation of transgenics via the random integration of exogenous DNA into the genome [22,34,36].

One bottleneck in performing functional genomics in *Hydractinia* and other cnidarians has been the inability to create heritable genetic modifications leading to true gene knockouts or knockins. With the advent of CRISPR/Cas9 genome editing, this goal is now within reach. To date, CRISPR/Cas9 has been used to disrupt genes in several cnidarians, including *Nematostella* [37–40], *Clytia* [41], *Acropora* [42], and *Hydractinia* [43]. These studies have each employed embryonic injection of one or more guide RNAs (sgRNAs) to disrupt a gene of interest, then evaluated phenotypes in developing embryos or adult animals derived directly from them. In addition, there has been one study in which a fluorescent reporter was knocked into a gene in *Nematostella [40].* Germline transmission of a CRISPR/Cas9 mediated genome edit has yet to be achieved in any cnidarian.

Here, we report the first use of CRISPR/Cas9 genome editing to generate knockins in *H. symbiolongicarpus.* Specifically, we use CRISPR/Cas9 to knock fluorescent reporters and small affinity tags into the endogenous the *Eukaryotic elongation factor 1 alpha* (*Eef1a*) locus. The transgenes were successfully transmitted through the germline to create fully transgenic animals and were stable over two generations of breeding. In addition, we demonstrate the utility of tagging the endogenous protein with two different affinity tags to perform *in vivo* and *ex vivo* protein studies. The ability to generate targeted gene knockins greatly increases the feasibility of studying the functions of cnidarian genes in their endogenous genomic environment, which can include complex *cis* and *trans* regulatory elements. It will also enable researchers to more efficiently create knockouts with fluorescent gene traps, to test hypotheses of protein function through targeted point mutations, and to engineer conditional transgenics that will allow gene expression to be controlled in site-, cell-, and/or time-specific ways.

## Results

### Identification of *Hydractinia Eef1a* and parental genotyping

Published and unpublished work from our laboratories has shown it is possible to create point mutations, small indels, and large deletions in the *Hydractinia* genome by injecting Cas9:sgRNA protein:RNA complexes into zygotes [43,44]. In the present study, we sought to test whether we could use CRISPR/Cas9 mediated homologous recombination to introduce exogenous DNA encoding fluorescent proteins into the *Hydractinia* genome. To test this, we focused on editing the *Eef1a* locus because it is a highly expressed housekeeping gene and would enhance our chances of visualizing the insertions.

To identify the *Hydractinia Eef1a* locus, we used previously published *H. symbiolongicarpus* RNA-Seq data [45] in conjunction with a draft genome assembly for a male strain, 291-10 [30]. From these mRNA and genomic sequences we identified two *Eef1a* alleles. Each allele encodes the same 465 amino acid peptide, spans ∼2.5 kb of genomic sequence, and is comprised of four exons (Fig 1A) (Sequences submitted to Genbank). The top hits resulting from BLASTP searches of these Eef1a peptides against the NCBI refseq_protein database were to Eef1a proteins from *Trichoplax adhaerens* (87% identity), *Branchiostoma floridae* (85% identity), and *Crassostrea virginica* (84% identity).

**Fig. 1.**
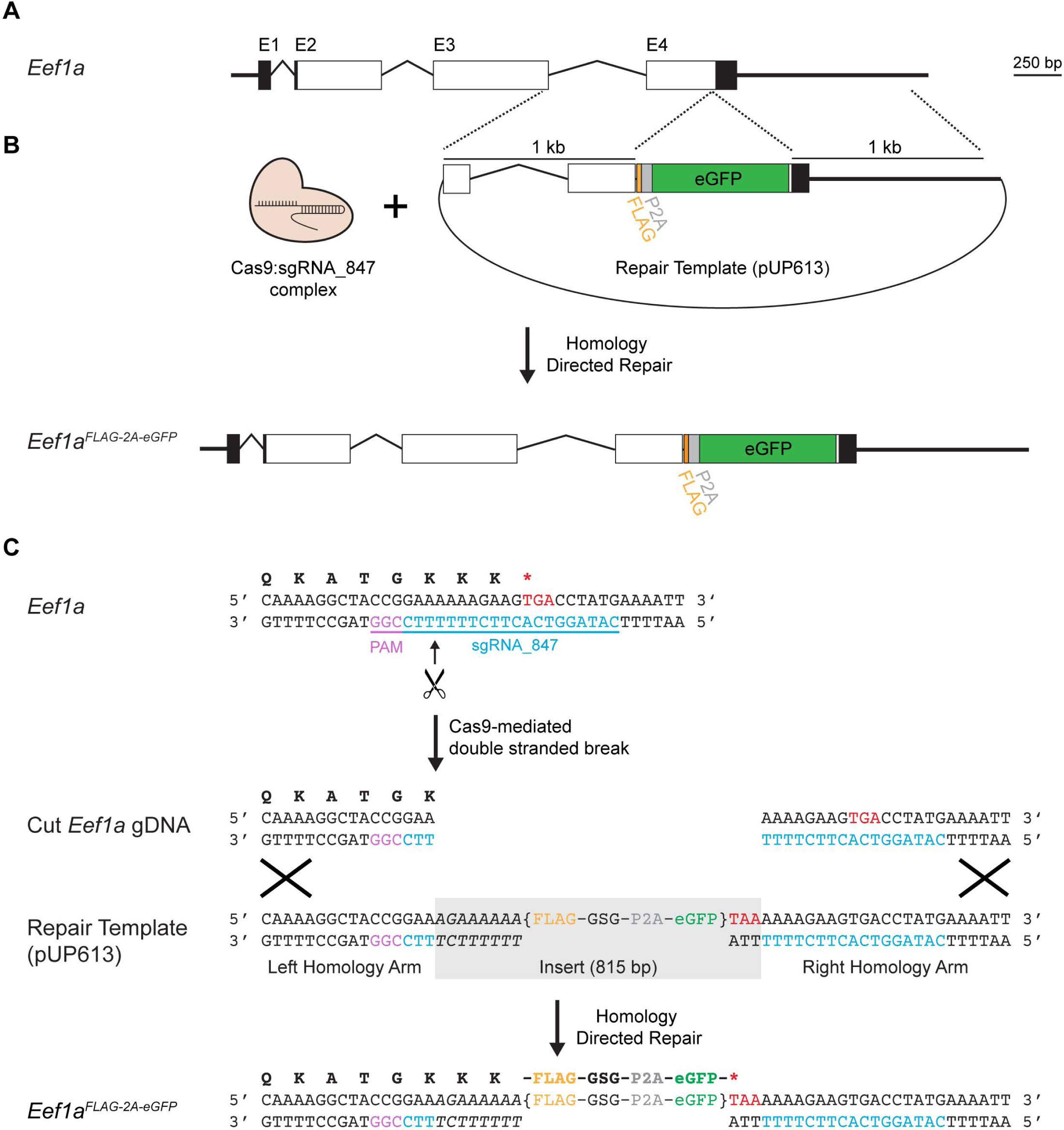
*Hydractinia Eef1a* gene structure and strategy for mutagenesis by homology directed repair. (A) Genomic organization of *Hydractinia Eef1a.* Boxes indicate exons, while angled lines indicate introns. Black areas are untranslated regions, white areas are coding sequences. The four exons are indicated as E1, E2, E3, and E4. (B) Overview of Cas9-mediated insertional mutagenesis. Injection of Cas9:sgRNA complexes and a plasmid containing a DNA repair template leads to the insertion of a FLAG-P2A-eGFP cassette via homology directed repair. (C) Nucleotide level view of insertional mutagenesis strategy. The single letter codes for the amino acid translation of the coding sequence is shown above the double-stranded DNA sequence. Bracketed text indicates the FLAG-P2A-eGFP cassette. Base pairs shown in italics were inserted to maintain the full Eef1a peptide.

Our microinjection experiments would ultimately require embryos, which we planned to generate by crossing 291-10 to a female colony, 295-8, which is a half-sibling of 291-10 (Additional file 1: Fig. S1). Because this female was likely to be heterozygous at *Eef1a*, we also genotyped it at the targeted region of *Eef1a.* We PCR-amplified the genomic region encoding exons 3-4 of *Eef1a*, which yielded a 2.0 kb and a 2.4 kb amplicon. Each amplicon was cloned and sequenced, revealing a ∼400 bp indel downstream of the *Eef1a* 3’UTR (Additional file 2). We concluded these sequences represented the two maternal *Eef1a* alleles. Aligning the four *Eef1a* alleles revealed several sequence variants but, as expected for a highly conserved housekeeping gene, identical coding sequences (Additional file 3).

### CRISPR/Cas9-mediated homologous recombination at the *Eef1a* locus

Our first goal was to test whether we could insert a transgene encoding an affinity tag and fluorescent reporter into the 3’ end of *Eef1a* via Cas9-mediated homology directed repair (Fig. 1B). To do this, we identified a NGG PAM site 14 bp upstream of the stop codon and used it to design a 20 bp sgRNA (sgRNA_847, Fig 1C) that was conserved across the four *Eef1a* alleles (Additional file 3). Local BLASTs of this sgRNA_847 sequence against the draft genome revealed no obvious off-target matches outside of *Eef1a* in the genome. We then created a plasmid to act as the repair template for homology-directed repair. This plasmid, pUP613 (Additional file 4), consisted of two 1,000 bp homology arms matching the sequences immediately before and after the predicted Cas9 cut site of one *Eef1a* allele from colony 291-10. Between these arms, we inserted 8 bp to complete the *Efla* coding sequence (italicized in Fig. 1C). Silent mutations in these 8 bp ensured the plasmid would not be targeted by Cas9:sgRNA_847 complexes. This was followed by sequences encoding a FLAG peptide, a Gly-Ser-Gly linker peptide, a P2A self-cleaving peptide, eGFP, and a stop codon, all codon-optimized for *Hydractinia* (Fig. 1C). When correctly inserted into the 3′ end of *Eef1a*, we predicted this construct would create a mutant allele, *Eef1a^FLAG-P2A-eGFP^*, resulting in the expression of a Eef1a-FLAG protein and a separate eGFP molecule that would act as a visual marker. Because the repair template did not include a promoter or splice acceptor site, untargeted integration of the plasmid into the genome would be unlikely to result in eGFP expression.

To test our construct, we injected zygotes with an injection cocktail containing 200 ng/µl repair template pUP613 and 1 µg/µl Cas9 complexed with 500 ng/µl sgRNA_847 (Fig2A). Pilot experiments comparing injected and non-injected embryos revealed an impact of injection on viability (Additional file 1: Fig. S2A). Despite this effect, we recovered 321 viable injected embryos (from three independent experiments), of which 14 (4.4%) were eGFP^+^ (Additional file 1: Fig. S2B). The location of eGFP^+^ expression ranged from several endodermal cells to long streaks of eGFP^+^ cells and remained present up to 144 hours post fertilization (hpf) (Fig. 2B). In control experiments in which we injected the repair template alone, we did not observe any eGFP^+^ cells (100 viable, injected embryos). A parallel set of experiments conducted with the same strains of *H. symbiolongicarpus* and the same injection cocktail in the Frank laboratory at the National University of Ireland, Galway also generated eGFP^+^ colonies.

**Fig. 2.**
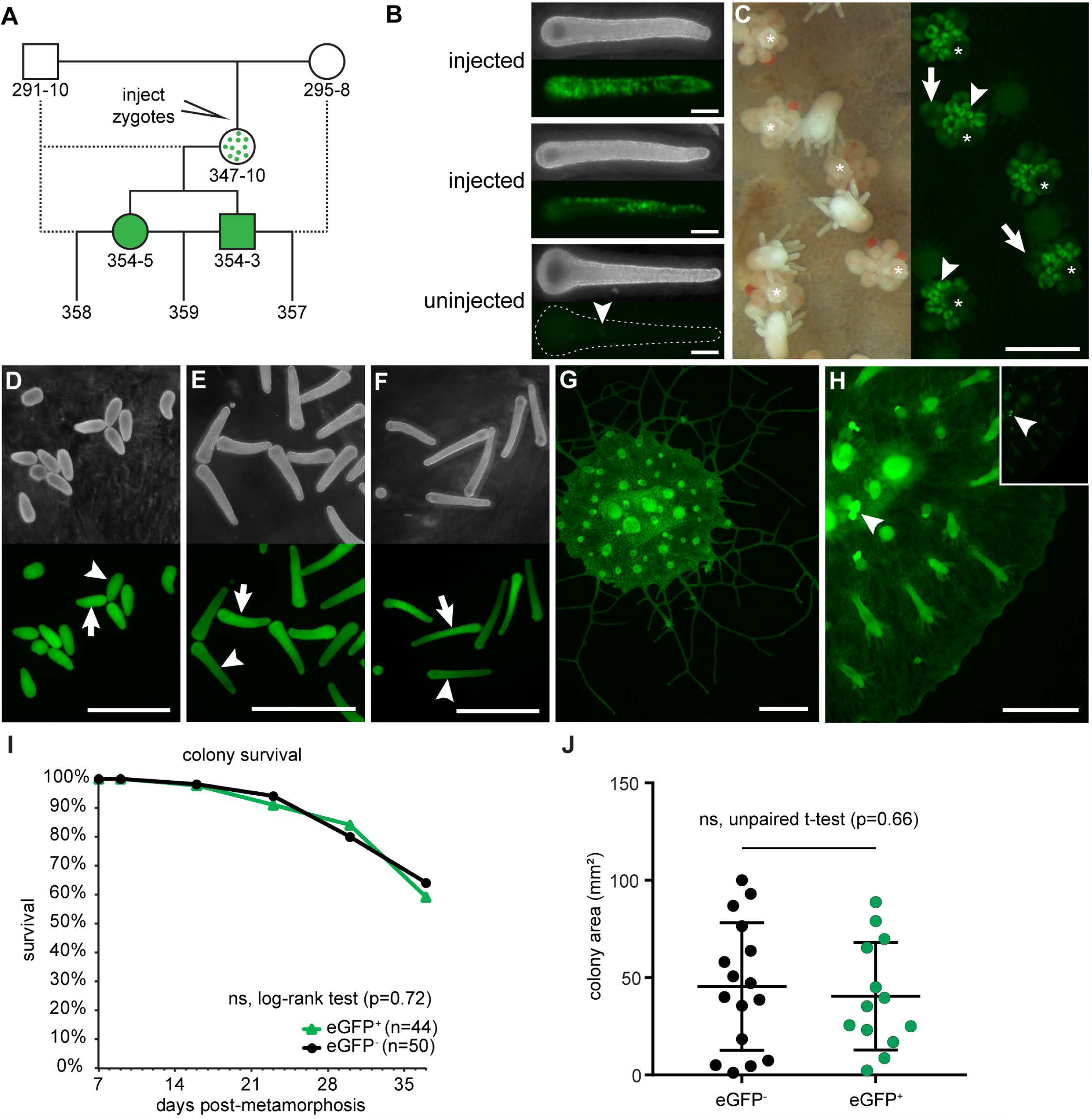
eGFP expression following Cas9 injection. (A) Breeding strategy and pedigree of eGFP^+^ colonies described in this paper. Colony 347-10 is a founder derived from an injected embryo and is assumed to be a mosaic of edited eGFP^+^ cells and wild-type eGFP^−^ cells. Dotted lines indicate backcrosses. (B) Larvae injected with Cas9:sgRNA complexes + repair template pUP613 at 144 hpf post-injection, showing mosaic expression of eGFP. An uninjected 144 hpf larvae is shown for comparison. Specks of green auto-fluorescence are common in uninjected larvae (arrowhead) but are easily distinguished from eGFP expression. (C) Left panel: dark-field image of colony 347-10, showing gonozooids (white asterisk). Right panel: Fluorescence micrograph of the same area of colony 347-10 showing eGFP expression. Developing oocytes (arrowheads) are intensely fluorescent, but become dim as they mature (arrows). (D) Darkfield (top) and fluorescence (bottom) micrograph of 24 hpf embryos from the backcross of 347-10 to 291-10, beginning to show slightly dimmer eGFP signal in some larvae (arrowhead) compared to others (arrow). (E) 72 hpf embryos from the same cross. eGFP^+^ (arrow) and eGFP^dim^ (arrowhead) larvae are now evident. (F) 168 hr-old embryos. The distinction between eGFP^+^ (arrow) and eGFP^dim^ (arrowhead) larvae is even more substantial. (G) Fluorescence micrograph of an explant of colony 354-3. (H) Fluorescence micrograph of an explant of colony 354-5. Image input and output levels were adjusted to reveal eGFP fluorescence in stolonal mat. Inset: Image with levels scaled automatically in NIS-elements, showing intense fluorescence of developing oocytes (arrowhead). (I) Survivorship of eGFP^+^ and eGFP^−^ colonies from population 357. (J) Mat area measurements of colonies from population 357. All scale bars = 1 mm.

We subsequently injected embryos and settled only those larvae with many eGFP^+^ cells (n=36). In all cases, we found that eGFP^+^ larvae metamorphosed into primary polyps with eGFP^+^ cells. As they developed into juvenile colonies, several of these animals lost eGFP expression. These were typically primary polyps that possessed few eGFP^+^ cells after metamorphosis. In 15 colonies, however, eGFP expression was stably maintained, albeit mosaic in most colonies.

In one eGFP^+^ colony, 347-10, we observed nearly ubiquitous eGFP expression at sexual maturity. In particular, we noted high levels of eGFP fluorescence in all developing oocytes (Fig 2C), raising the possibility that the *Eef1a^FLAG-P2A-eGFP^* allele might be present in the germline. To test this, we backcrossed colony 347-10 to its male parent (Fig. 2A). At 24 hpf, 100% of larvae obtained from this cross were eGFP^+^ (Fig. 2D), although some appeared slightly dimmer than others. By 72 hpf, we could clearly distinguish eGFP^+^ from eGFP^dim^ larvae (Fig. 2E), suggesting that the eGFP^+^ had received a germline transmitted *Eef1a^FLAG-P2A-eGFP^* allele, while eGFP^dim^ fluoresced due to maternal loading of either eGFP mRNA or protein in the oocyte. This low level of eGFP signal remained in the larvae up to seven days post-fertilization, while eGFP+ larvae accumulated bright eGFP signal, particularly in the endodermal cells (Fig. 2F). eGFP^+^ larvae that we settled developed into eGFP^+^ adults with high levels of eGFP expression, while eGFP^dim^ larvae developed into colonies without eGFP expression.

To further assess the stability of this transgene, we grew several of these eGFP^+^ colonies to sexual maturity and never observed loss of eGFP expression. We then selected a male and a female (Figs. 2G and 2H, respectively), backcrossed them to their parents, and crossed them to each other (Fig. 2A). At 24 hpf, larvae resulting from the backcross of the male eGFP^+^ colony could readily classified as eGFP^+^ or eGFP^−^ (not shown). In contrast, larvae resulting from both crosses involving the female eGFP^+^ colony displayed either high or low eGFP signal similar to that observed coming from the founder colony, 347-10. This phenotype resolved into eGFP^+^ and eGFP^dim^ classes by 72 hpf. At 96 hpf, we counted the number of eGFP^+^ and eGFP^dim^ or eGFP^−^ larvae in each cross and confirmed that the inheritance pattern of eGFP expression was consistent with mendelian inheritance of a single *Eef1a^FLAG-P2A-eGFP^* allele from colonies 354-3 and 354-5 (Table 1).

**Table 1.** Mendelian inheritance of eGFP expression in testcrosses

| Cross<sup>a</sup> | Parents | Hypothesized Genotypes |  | N<sup>c</sup> | Phenotype<sup>d</sup><br>eGFP<sup>+</sup>/eGFP<sup>-</sup> | Goodness of fit |
| --- | --- | --- | --- | --- | --- | --- |
|  |  | Parental<br>(male x female) | Offspring<sup>b</sup> |  |  |  |
| 357 | 354-3<br>x<br>295-8 | *Eef1a*<sup>FLAG-P2A-eGFP/+</sup><br>x<br>*Eef1a*<sup>+/+</sup> | *Eef1a*<sup>FLAG-P2A-eGFP/+</sup><br>*Eef1a*<sup>+/+</sup> (1:1) | 263 | 136/127 | $\chi^2 = 0.647$,<br>$p=0.58$ |
| 358 | 291-10<br>x<br>354-5 | *Eef1a*<sup>+/+</sup><br>x<br>*Eef1a*<sup>FLAG-P2A-eGFP/+</sup> | *Eef1a*<sup>FLAG-P2A-eGFP/+</sup><br>*Eef1a*<sup>+/+</sup> (1:1) | 124 | 71/53 | $\chi^2 = 0.018$,<br>$p=0.11$ |
| 359 | 354-3<br>x<br>354-5 | *Eef1a*<sup>FLAG-P2A-eGFP/+</sup><br>x<br>*Eef1a*<sup>FLAG-P2A-eGFP/+</sup> | *Eef1a*<sup>FLAG-P2A-eGFP/FLAG-P2A-eGFP</sup><br>*Eef1a*<sup>FLAG-P2A-eGFP/+</sup><br>*Eef1a*<sup>+/+</sup> (1:2:1) | 111 | 81/30 | $\chi^2 = 0.777$,<br>$p=0.62$ |

<sup>a</sup>As described in Fig 2A; <sup>b</sup>Expected ratio is indicated in parentheses. <sup>c</sup>larvae at 96 hpf; <sup>d</sup>eGFP<sup>dim</sup> larvae were scored as eGFP<sup>-</sup> in crosses 358 and 359.

To test for potentially deleterious fitness effects of the *Eef1a^FLAG-P2A-eGFP^* allele, we settled 44 eGFP^+^ and 50 eGFP^−^ larvae from the 357 cross and followed them for four weeks. We observed no difference in survival or total area of the mat (a proxy for colony size) between the two groups (Figs. 2I-J). To date, we have continued to culture five animals from the 357 population for seven months and observed stable eGFP expression in each one.

To determine whether we had correctly inserted the FLAG-P2A-eGFP cassette into the *Eef1a* locus, we designed PCR primers to amplify the transgene from eGFP^+^ animals. Repeated attempts to amplify the full 2921 bp cassette from eGFP^+^ colonies with primers Pr845 and Pr846 (Fig. 3A) were unsuccessful. We concluded this was likely due to preferential amplification of the shorter, unedited *Eef1a* allele. We therefore developed a strategy to amplify each side of the inserted transgene. To avoid amplifying repair template DNA that may have integrated elsewhere in the genome, we designed each primer pair such that one primer annealed to the endogenous *Eef1a* locus outside the region matching the homology arms in pUP613 and the other to the insert (Fig. 3A and Additional file 1: Table S1). Using primers to the 5’ homology arm, we were able to amplify a band of the expected size from the founder (347-10) and its eGFP^+^ progeny, but not their eGFP^−^ siblings (Fig. 3B). We cloned and sequenced these amplicons and found that they matched the expected sequence of the *Eef1a^FLAG-P2A-eGFP^* allele (Additional file 5). Using primers flanking the 3’ homology arm, we were also able to amplify products from eGFP^+^ colonies, but these were smaller than expected. Cloning and sequencing them revealed that they matched expected sequence of the *Eef1a^FLAG-P2A-eGFP^* allele except for a 681 bp deletion downstream of the *Eef1a* 3’ UTR. These experiments confirmed that colonies 354-3 and 354-5 inherited the same *Eef1a^FLAG-P2A-eGFP^* allele from the founder colony, 347-10, but do not entirely rule out the possibility that 347-10 was a mosaic possessing multiple edited *Eef1a* alleles. Taken together, these results show that CRISPR/Cas9-mediated homologous recombination can be used to introduce exogenous DNA into target sites in the *Hydractinia* genome, and that these sequences can be stably inherited over multiple generations.

**Fig. 3.**
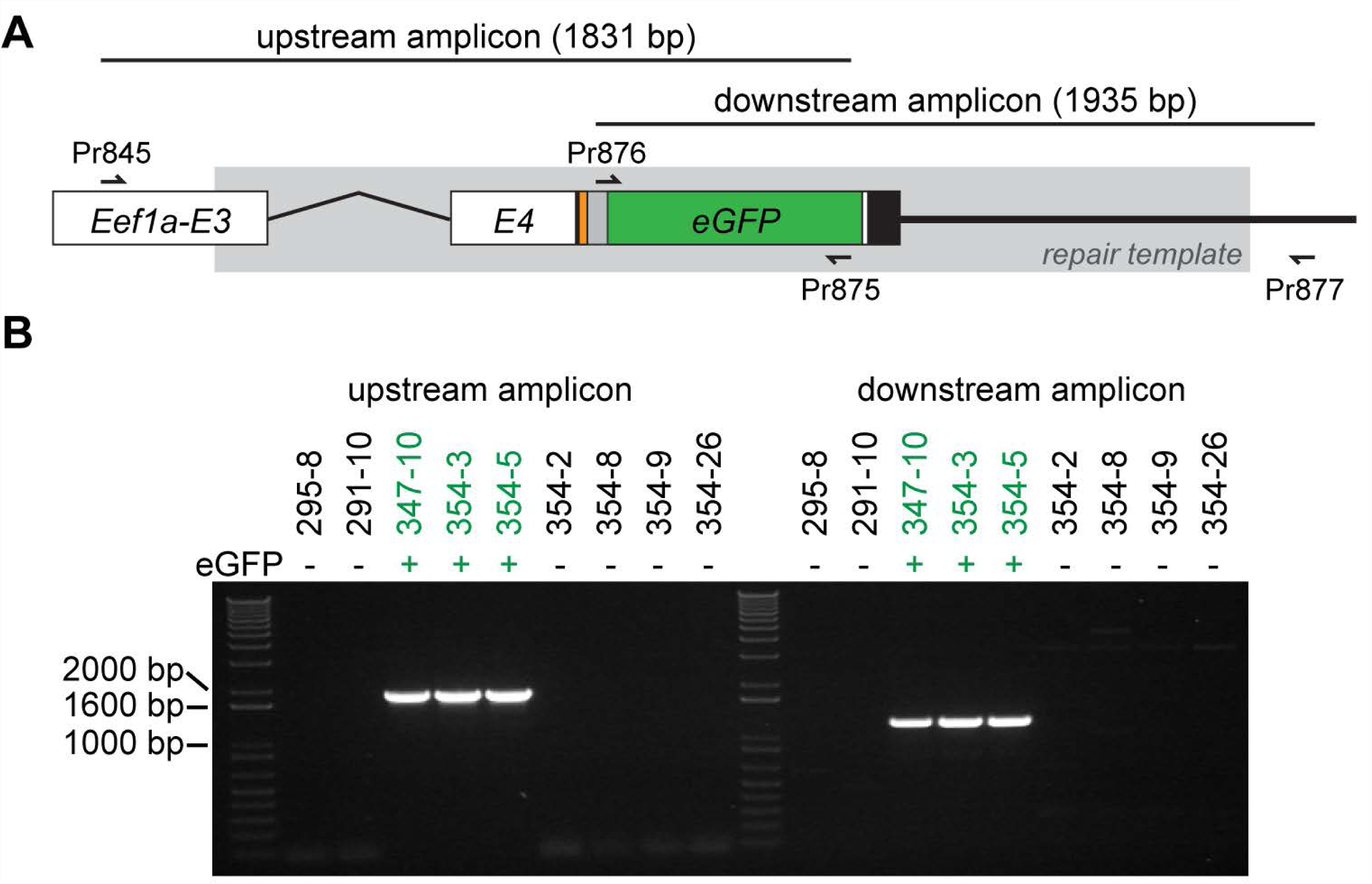
Validation of homologous recombination at the *Eef1a* locus. (A) Schematic of the expected sequence for the edited region of *Eef1a^FLAG-P2A-eGFP^.* Exons are abbreviated E3 and E4. Sequence matching the repair template is shown in gray box. PCR primers are shown as small arrows. Coding sequence is colored as in Fig. 1. (B) Amplification of upstream and downstream regions of *Eef1a^FLAG-P2A-eGFP^* from eGFP^+^ and eGFP^−^ colonies.

### The FLAG octopeptide enables specific tagging of endogenous proteins in *Hydractinia*

One potential use for CRISPR/Cas9 genome editing is to tag endogenous proteins of interest with short epitopes to facilitate *in vivo* and *in vitro* protein detection via immunolocalization and western blotting. This strategy could obviate the laborious task of creating and validating antibodies to target proteins for such studies. An additional advantage of using small epitope tags instead of fluorescent fusion proteins is that short tags are less likely to interfere with protein stability and function. To test whether we can specifically detect FLAG-tagged proteins in *Hydractinia* using commercially available reagents, we isolated total protein from transgenic and nontransgenic colonies and probed them with an anti-FLAG monoclonal antibody (M2) in western blots (Fig. 4A). We observed a strong, specific signal in colonies expressing the Eef1a-FLAG protein, with no detectable background in colonies expressing only untagged Eef1a. The absence of additional bands at the expected size of the intact Eef1a-FLAG-P2A-eGFP protein (80.8 kDa), even after 60 minutes of exposure to film, also demonstrated that the P2A self-cleaving peptide can efficiently separate two peptides in vivo in *Hydractinia.*

**Fig. 4.**
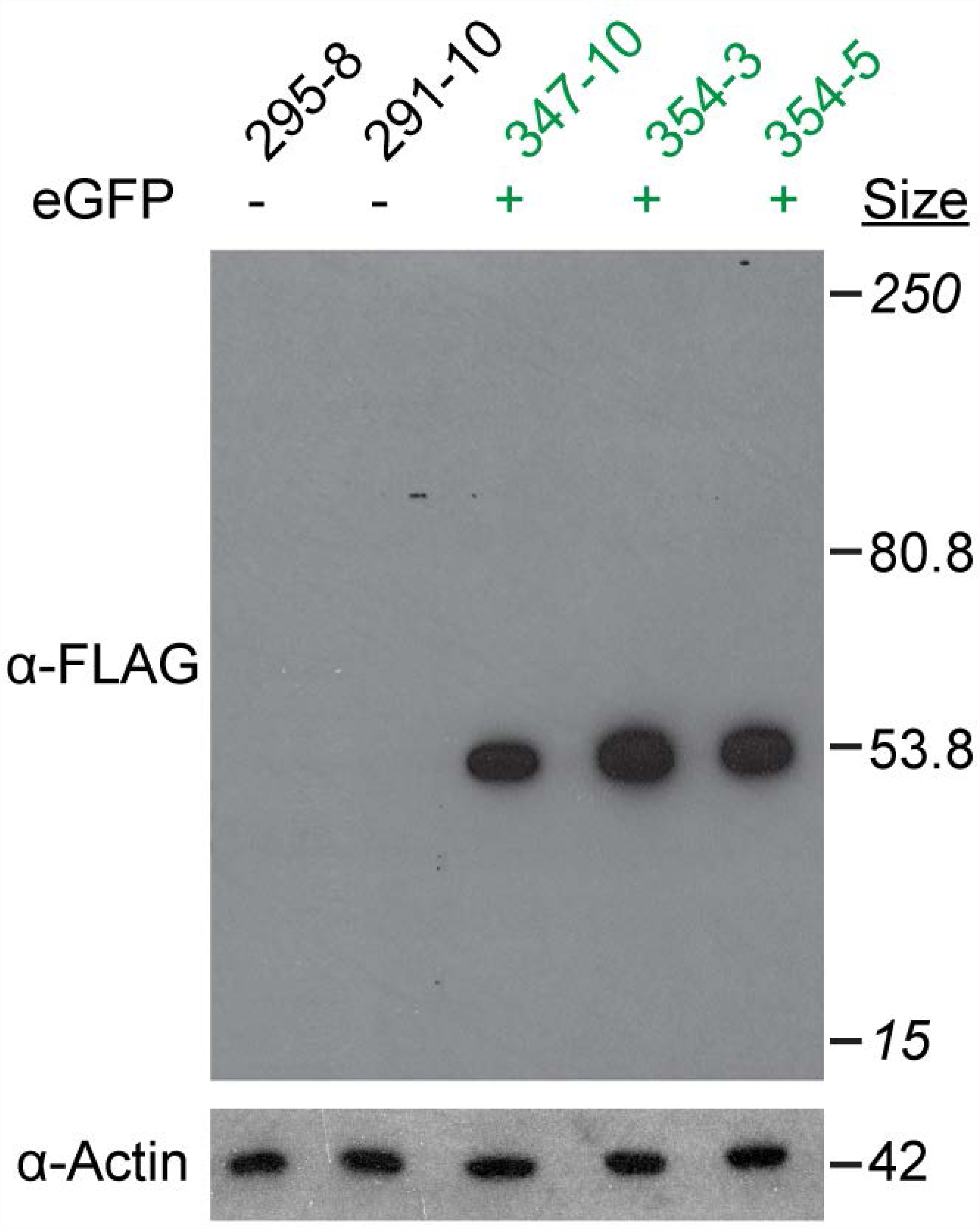
Specificity of the FLAG octopeptide in *Hydractinia*. Western blot of total protein lysates from *Eef1a^FLAG-P2A-eGFP^* transgenic and nontransgenic colonies. Sizes in italics are the top and bottom bands of the molecular size marker.

To test whether the FLAG octopeptide would be suitable for immunolocalizations, we probed whole mounts of transgenic and non-transgenic gastrozooids with the same anti-FLAG antibodies. We observed strong but diffuse staining in the transgenic polyps (Figs. 5A-C and Additional files 6-7), which was consistent with ubiquitous cytoplasmic expression of Eef1a-FLAG. In addition, we noted cells with large nuclei and a polygonal morphology sparsely distributed within the body column (Figs. 5C-D). The morphology of these cells was similar to that reported for *Hydractinia* interstitial stem cells (i-cells) [e.g.,22]. In contrast, we observed very little signal from the non-transgenic control (Figs. 5E-G and Additional files 8-9). This signal was dramatically reduced in secondary-only controls (Additional file 1: Fig. S3), indicating it was likely the result of minimal non-specific staining from the FLAG antibody. These results suggest that tagging endogenous proteins with FLAG epitope tags could be a viable strategy for immunolocalization, provided appropriate controls are performed.

**Fig. 5.**
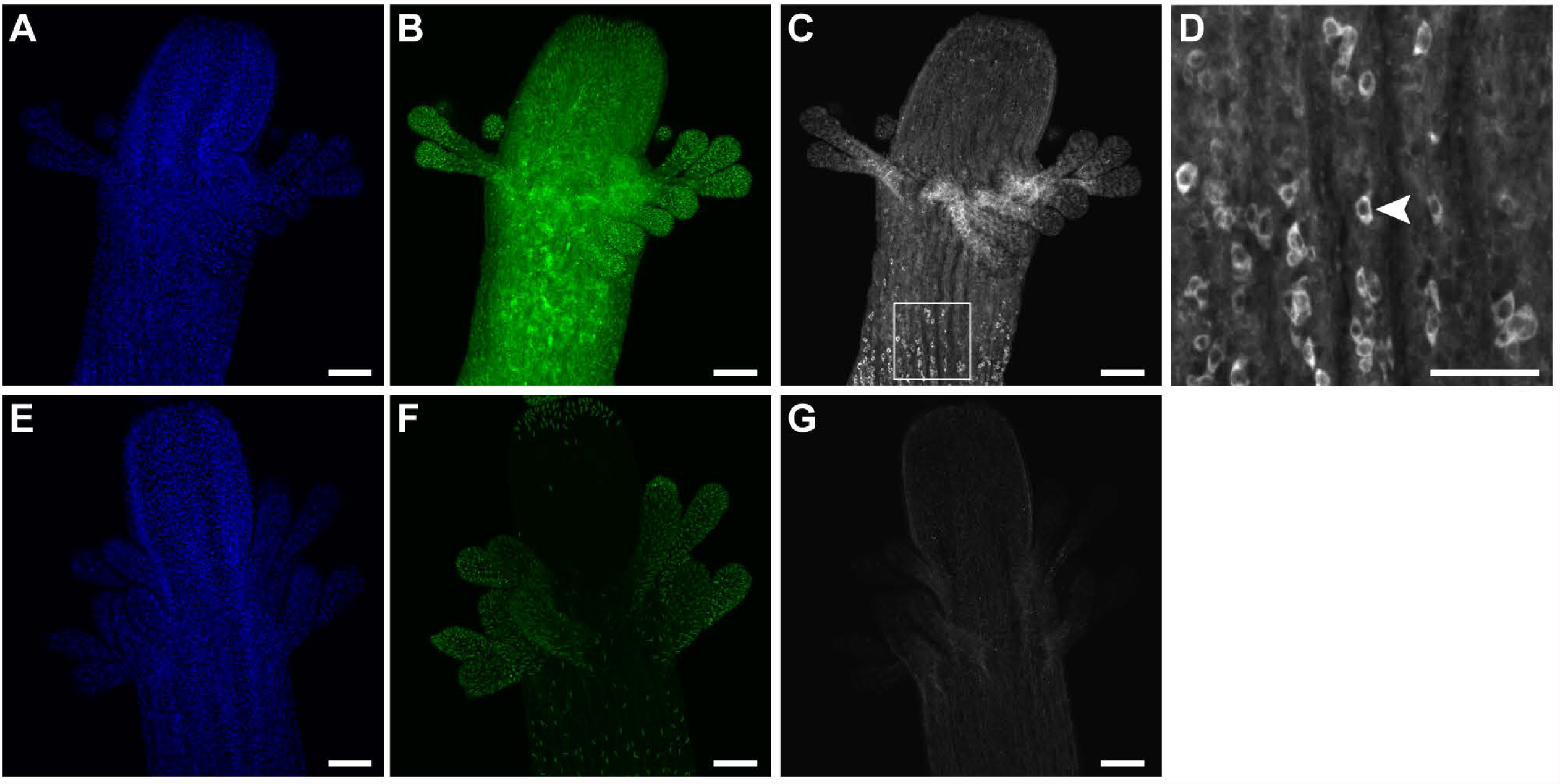
Immunolocalization of Eef1a-FLAG. (A-D) *Eef1a^FLAG-P2A-eGFP^* transgenic gastrozooids fixed, permeabilized, and stained with anti-FLAG primary antibodies and Alexa 647 conjugated secondary antibodies, followed by DAPI, and imaged on a laser scanning confocal microscope. (A,B) DAPI staining and eGFP expression in the transgenic gastrozooid, respectively. (C) Anti-FLAG staining in the transgenic gastrozooid, showing strong, diffuse signal throughout with sparse, intensely stained cells. (D) An inset of panel C showing intensely stained cells with polygon-like morphology (e.g., arrowhead). (E-H) Nontransgenic controls imaged and analyzed with identical settings. (E,F) DAPI staining and green fluorescence, respectively. Nematocyst capsules exhibit bright autofluorescence. (G) Anti-FLAG staining, showing faint background staining. Scale bars =100 µm, except for (D), which is 50 µm.

### Germline transmitted *Eef1a^FLAG-P2A-eGFP^* is expressed without evidence of variegated expression

Because *Eef1a* is a constitutively expressed housekeeping gene, we expected to observe eGFP expression in all cells at all developmental stages. However, previous work has shown that germline transmitted, randomly integrated transgenes driven by promoters of housekeeping genes can display an expression pattern in which cells appear to either stochastically express or not express the transgene [36]. The mechanism responsible for this variegated expression pattern is unknown, but for transgenes driven by actin promoters, it can persist even after two generations of germline transmission (ML Nicotra, unpublished data).

To determine whether the germline transmitted *Eef1a^FLAG-P2A-eGFP^* allele would exhibit similar variegated expression, we investigated its expression in detail. Multiphoton imaging of intact gastrozooids, gonozooids, stolons, and mat edges revealed no evidence of variegated expression in either the epidermal or gastrodermal cell layers (Figs. 6A-G, J, K and Additional files 10-15). The intensity of eGFP signal did not allow us to determine whether expression was uniform across all cell types. We also investigated 120 hpf eGFP^+^ larvae from the 357 cross and found robust eGFP expression in all ectodermal and endodermal cells (Figs. 6H-I and Additional files 16-17). We therefore concluded that, as expected, the *Eef1a^FLAG-P2A-eGFP^* transgene is at least expressed in all epithelial cells, with no detectable evidence of variegated expression.

**Fig. 6.**
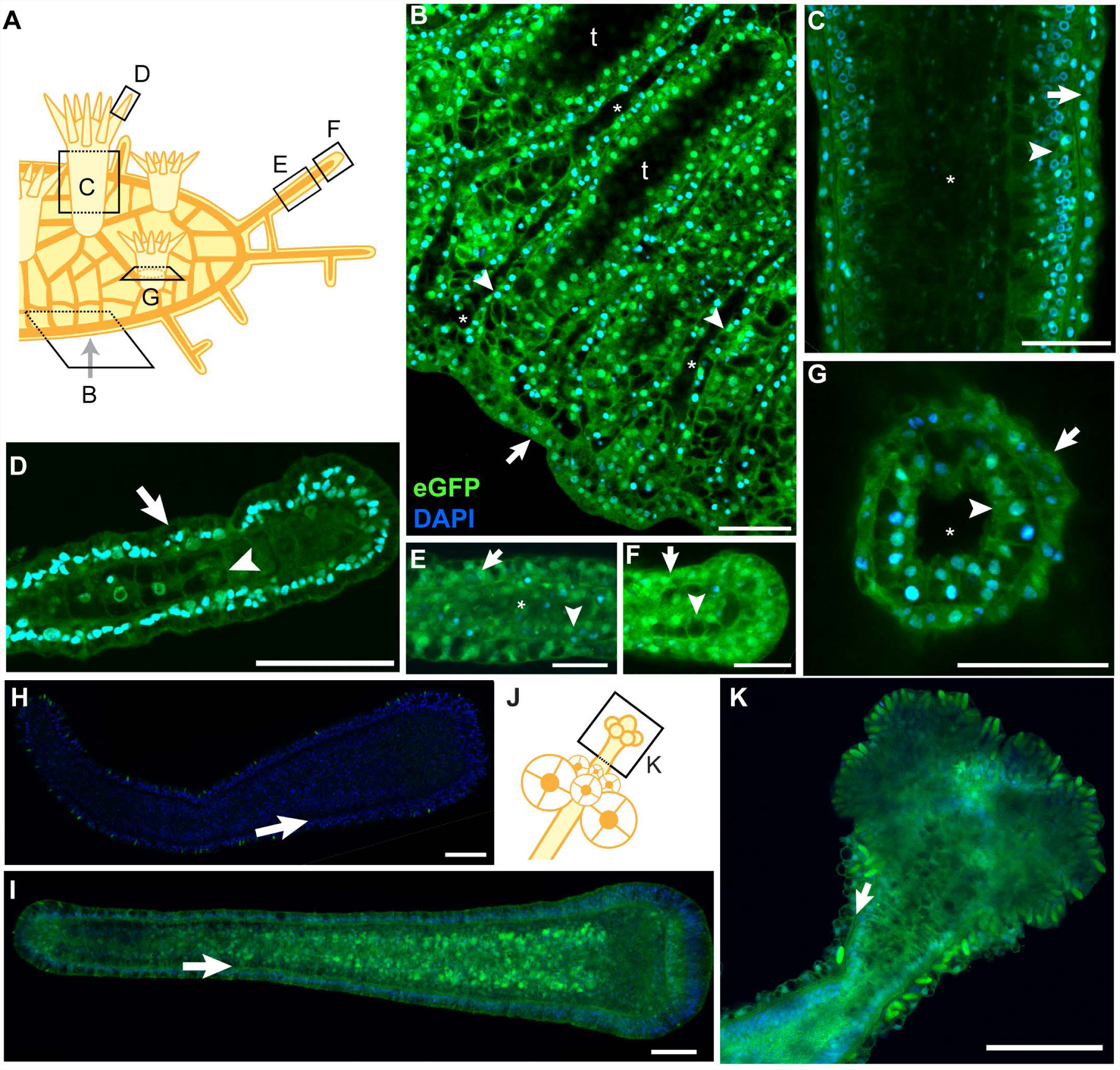
No evidence of variegated expression of the *Eef1a^FLAG-P2A-eGFP^* transgene in epithelial cell layers. (A) Schematic of colony showing orientation of multiphoton micrographs in B-G. (B-G) Multiphoton micrographs of *Eef1a^FLAG-P2A-eGFP^* expressing colonies stained with DAPI. Epidermal cell layers are indicated with arrows, while gastrodermal cell layers are indicated with arrowheads. (B) Edge of stolonal mat. Focal plane is through the center of the gastrovascular canals, leaving areas of the stolonal mat lacking canals dark because they are below the plane of focus (“t”). Asterisks denote lumens of gastrovascular canals. (C) Body of gastrozooid. Asterisk indicates lumen. (D) End of tentacle. (E) Tip of stolon. (F) Middle of stolon. Asterisk denotes stolon lumen. The lumen has collapsed toward the bottom of the image. (G) Base of polyp imaged through bottom of colony. Asterisk indicates lumen. (H) 120 hpf eGFP- larva from the 357 cross. Arrow denotes border of ectodermal and endodermal cell layers. (I) 120 hpf eGFP+ larva from the 357 population. Arrow same as in H. (J) Schematic of female gonozooid, showing plane of image in (K). (K) Confocal image of head of female gonozooid. Arrow denotes border of epidermal and gastrodermal cell layers. All scale bars = 50 microns.

### *In vivo* expression and function of alternative affinity tags and fluorescent reporters

While the FLAG epitope was effective for immunoblotting and immunolocalization studies, repeated attempts to immunoprecipitate the tagged Eef1a protein failed. Therefore, we decided to test the StrepII-FLAG tandem affinity purification (SF-TAP) tag (Fig. 7A) [46]. The SF-TAP tag is composed of a doublet Strep II tag followed by a single FLAG peptide and was chosen because it would potentially enable efficient purification of tagged proteins and their interaction partners via tandem affinity purification [46]. In addition, we replaced the eGFP with tdTomato because it is one of the brightest fluorescent proteins available [47] and can be used in combination with eGFP during imaging. The resulting repair template (pUP660; Additional file 18) was expected to create an Eef1a-SF peptide and a separate tdTomato peptide.

**Fig. 7.**
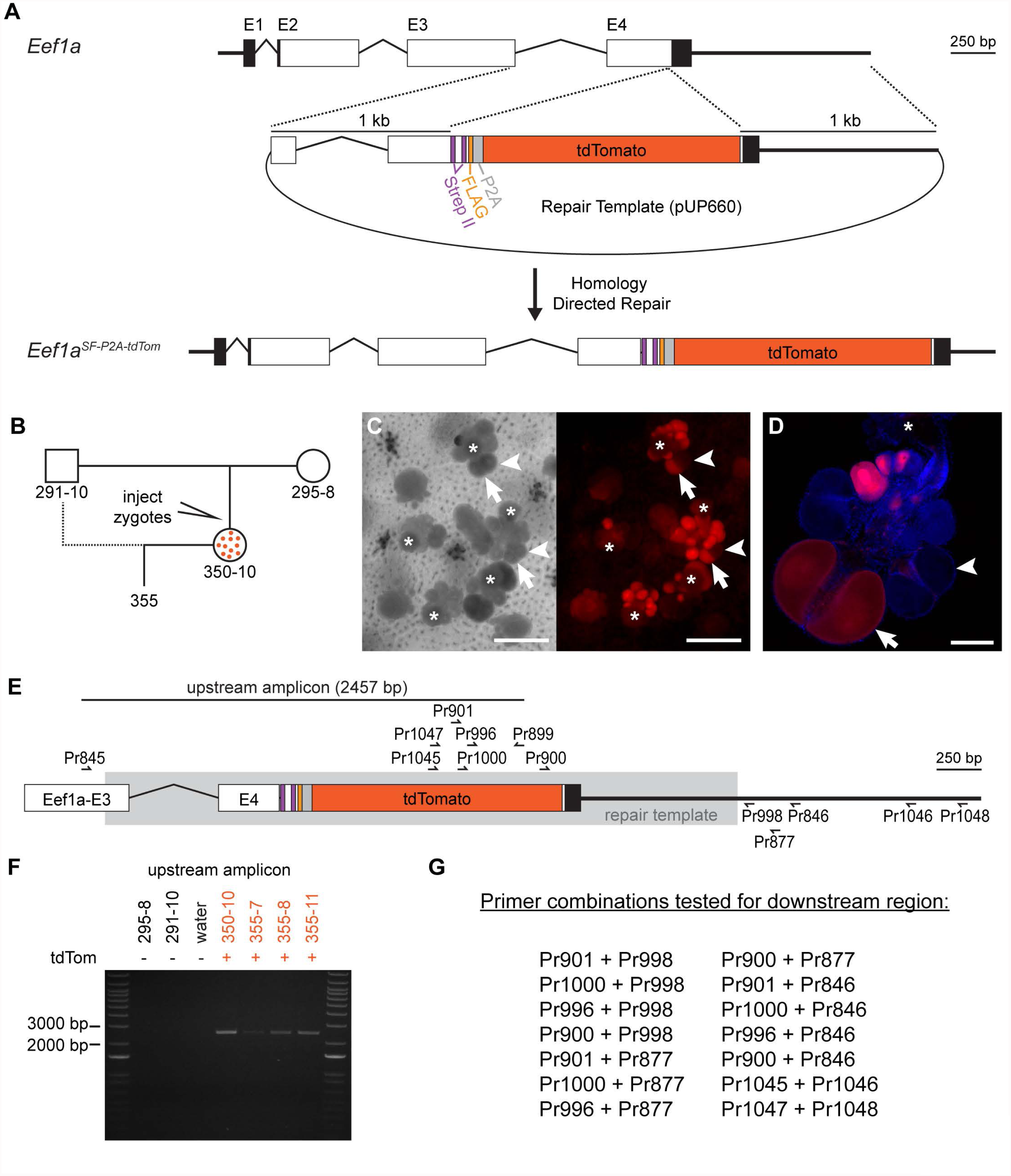
Creation of the *Eef1a*^SF-P2A-tdTom^ allele. (A) Overview of Cas9-mediated insertional mutagenesis. Injection of Cas9:sgRNA complexes and a plasmid containing a DNA repair template leads to the insertion of a SF-P2A-tdTomato cassette via homology directed repair. (B) Pedigree of tdTomato^+^ colonies in this paper. Colony 350-10 is a founder derived from an injected embryo and is a mosaic of edited tdTomato^+^ cells and wild-type tdTomato^−^ cells. (C) Left panel: light transmitted light micrograph of colony 350-10, showing gonozooids (white asterisk indicates the head). Arrows and arrowheads point to the same eggs as in the right panel. Right panel: Fluorescence micrograph of the same area of colony 350-10 showing tdTomato^+^ eggs. Single gonozooids contain oocytes that are fluorescent (arrow) and non-fluorescent (arrowheads). Scale bars = 1 mm. (D) Confocal image of one gonozooid, showing mixture of tdTomato positive and negative eggs. Scale bar = 100 µm. (E) PCR primers used to validate insertion at the *Eef1a* locus. Region of 100% identity with the repair template is shown in gray box. (F) PCR amplification of upstream region from parental and tdTomato^+^ colonies. (G) Primer combinations tested in order to amplify the downstream region of *Eef1a^SF-P2A-tdTom^* allele.

We co-injected pUP660 with Cas9:sgRNA_847 complexes into zygotes from parents 291-10 and 295-8 (Fig. 7B). Similar to previous experiments, we observed tdTomato^+^ cells in a portion of the injected embryos at 72 hpf and selected only those larvae with high levels of tdTomato expression for metamorphosis into colonies. Of 38 such colonies, one colony, 350-10, developed tdTomato^+^ eggs. In contrast to colony 347-10, however, not all eggs were fluorescent, even within the same gonophore (Figs. 7C-D). We backcrossed this colony to 291-10 and obtained several tdTomato^+^ colonies.

To validate correct insertion of the SF-P2A-tdTomato cassette, we attempted to amplify regions upstream and downstream of the cassette using a similar PCR-based strategy to that employed for *Eef1a*^FhAG-F2A-eGFP^ (Fig. 7E). For the upstream region, we obtained bands of the expected size from tdTomato^+^ colonies (Fig. 7F), which we cloned and sequenced and found identical to the expected sequence of the edited *Eef1a^SF-P2A-tdTom^* allele (Additional file 19). In contrast, we were unable to amplify the downstream region, despite testing 11 PCR primers in 14 pairwise combinations (Fig 7G). These results led us to conclude we had successfully created the *Eef1a^SF-P2A-tdTom^* mutant allele and, since tdTomato was functional, that the sequence was intact at least through the stop codon. We could not, however, confirm the sequence of the downstream genomic region.

In culturing colonies from the 355 population, we also observed a marked fitness defect associated with the *Eef1a^SF-P2A-tdTom^* allele. Whereas tdTomato^−^ larvae metamorphosed and grew into robust colonies, their tdTomato^+^ siblings required meticulous hand-feeding for several weeks, died at a high rate, and grew slowly. Despite this defect, we were able to obtain enough tissue to test the suitability of the SF-TAP tag for *in vivo* protein studies.

We first tested whether we could detect both tags via in western blots on total cell lysates of tdTomato^+^ and tdTomato^−^ colonies. Using anti-FLAG antibodies, we observed a band of the expected molecular weight for an Eef1a protein bearing a C-terminal SF-TAP tag in an animal bearing a germline transmitted allele, but no signal in the mosaic founder (Fig. 8A). Using an anti-Strep II monoclonal antibody, we similarly observed a strong signal from the fully transgenic colony and no signal from the mosaic founder. (Fig. 8B). After over-exposure on film, however, a faint band appeared for the founder, likely reflecting the small number of properly edited cells in this colony (Fig. 8B). In addition, we observed a 15 kDa band in all colonies, suggesting this antibody cross-reacts with some variants of an endogenous *Hydractinia* protein. These results further confirmed we had, indeed, created the *Eef1a^SF-P2A-tdTom^* allele, despite our inability to amplify the downstream genomic region via PCR.

**Fig. 8.**
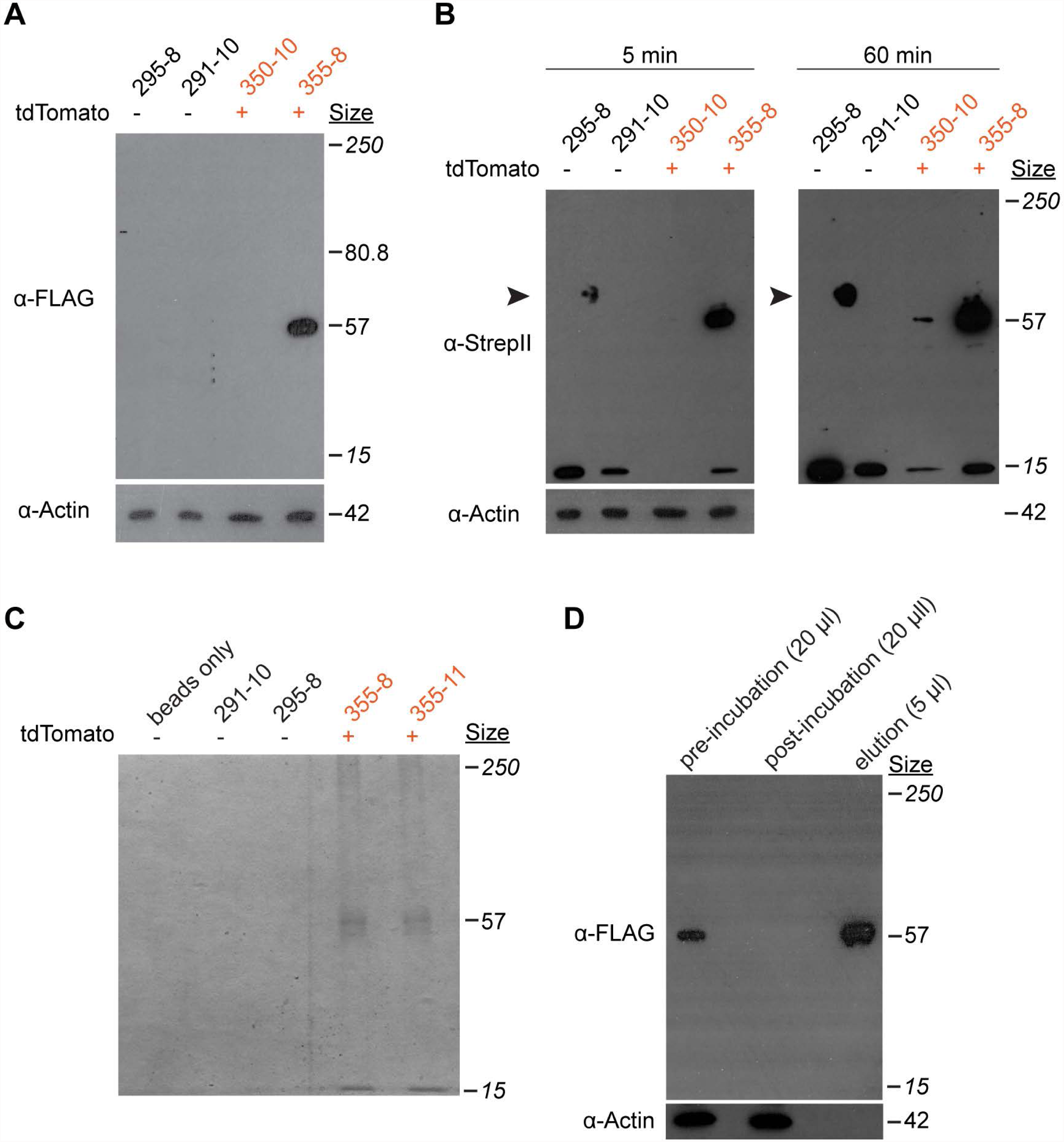
Use of the StrepII-FLAG tag in immunoblots and pull-downs. (A) Western blot for FLAG on total protein lysates *Eef1a^FLAG-P2A-tdTom^* transgenic and nontransgenic colonies. Sizes in italics are the top and bottom bands of the molecular size marker. (B) Western blot of Strep tag on total protein lysates from the same colonies. The blot was treated to induce chemiluminescence and exposed to film for 5 minutes (left), followed by a second exposure for 60 minutes (right). The 15 kDa band suggests this antibody cross-reacts with some variants of an endogenous *Hydractinia* protein. Arrowhead indicated non-specific staining of protein ladder. (C) Coomassie stained SDS-PAGE gel of elutions from streptavidin coated beads following incubation in total protein lysates from the indicated colonies. (D) Western blot against the FLAG peptide on the indicated amount of total protein lysate from colony 355-8, before and after incubation with the anti-streptavidin beads. The elution from the beads was run alongside for comparison.

Finally, to test whether we could use the SF-TAP tag to purify proteins from the *Hydractinia* proteome, we harvested tissue from transgenic and non-transgenic colonies, prepared whole cell lysates, incubated them with Streptactin conjugated agarose beads, and eluted the bound protein. SDS-PAGE of the elutions followed by coomassie staining revealed band of the expected size for Eef1a-SF from transgenic but not control colonies. We also observed a faint band smaller than the expected band, which may be a protein that specifically or non-specifically binds to Eef1a. To test the efficiency of the pull-down, we also probed whole cell lysates before and after the pull down with anti-FLAG antibodies and observed a complete loss of FLAG signal, indicating complete capture of the Eef1a-SF protein. These results demonstrate the suitability of the Strep II tag to purify in vivo tagged proteins from *Hydractinia.*

## Discussion

Our study demonstrates that it is possible target the insertion of exogenous DNA into the *Hydractinia* genome via CRISPR/Cas9 genome editing. Specifically, we inserted either a FLAG-P2A-eGFP or a SF-P2A-tdTomato coding sequence into the *Eef1a* locus. With moderate effort, we were able to generate mosaic founders that transmitted each transgene through the germline, thus generating fully transgenic animals. We then used these animals to confirm the correct insertion of the reporter construct. Transgene expression in these offspring was stable throughout the duration of our experiments and across two generations of breeding. With this work, *Hydractinia* becomes the first cnidarian to display successful germline transmission of CRISPR/Cas9 generated transgenes and, along with *Nematostella* [40]. only the second cnidarian in which CRISPR/Cas9 mediated homologous recombination has been achieved.

In both cases of germline transmission in *Hydractinia*, we observed that the genomic regions upstream of the inserted transgene matched the expected sequence, but downstream regions did not. Specifically, we discovered a 681 bp deletion in the *Eef1a^FLAG-P2A-eGFP^* allele and were unable to amplify the downstream region of the *Eef1a^FLAG-P2A-tdTom^* allele. One interpretation of these results is that homology directed repair (HDR) in *Hydractinia* could be more error-prone than anticipated. Specifically, our results are consistent with a scenario in which the lesion created by Cas9 results in significant alterations to both up- and downstream regions, which are then repaired by HDR or another form of DNA repair. Since we used fluorescence to screen for founder colonies, we were only able to detect transgenic animals in which the upstream regions preserved the reading frame of the reporter. Although the presence of downstream mutations did not create a problem in this study, it could complicate future work that depends on the sequence on both sides of an inserted transgene. It will therefore be important to determine the conditions that promote and/or prevent these mutations.

One variable that could affect the prevalence of mutations in surrounding regions is the size of the homology arms in the repair template. In this study, we followed the example of Ikmi *et al.* [40] and used 1kb homology arms. However, homology arms of ∼500 bp are routinely used to insert 1kb transgenes in mammalian cells, while shorter transgenes can be inserted using homology arms as short as 40 bp and a single stranded oligoDNA template [48]. Shortening the homology arms in *Hydractinia* might narrow the region available for homologous recombination and thus decrease the odds of recovering animals with large mutations. Further investigating the effect of homology arm length on the efficiency and fidelity of CRISPR/Cas9 mediated HDR in *Hydractinia* will be useful.

Our results demonstrate the feasibility of using short affinity tags to track endogenous proteins *in vivo.* One immediate application is to tag proteins in order to study their expression via immunolocalization or western blot. We showed here that the FLAG octopeptide is suitable for both applications in *Hydractinia.* This approach could, in some instances, eliminate the need to generate and validate antibodies in order to study a protein of interest. Another potential application is the purification of *in vivo* expressed proteins to study biochemical features such as post-translational modifications or interaction partners. Using relatively small amounts of tissue (area ∼ 6 cm^2^) and a simple incubation procedure with Streptactin beads, we were able to extract all detectable StrepII-tagged Eef1a protein from total cell lysates in sufficient amounts to visualize on a coomassie stained polyacrylamide gel. One caveat to this result is that Eef1a is typically one of the most abundant proteins in eukaryotic cells, making it an easy target. Nonetheless, we expect that purifying less abundant proteins will be as simple as increasing the amount of starting tissue, a straightforward task in a clonally reproducing animal like *Hydractinia.* In short, this approach should enable *Hydractinia* researchers to study previously intractable questions regarding protein biochemistry using commercially available reagents.

To the best of our ability, we have sought to determine whether every cell in a transgenic colony is marked with a fluorescent reporter. In no instance did we see evidence of cells lacking expression of either eGFP or tdTomato when it is placed under control of the endogenous *Eef1a* promoter. This suggests that *Eef1a* will be a useful landing site for the ubiquitous expression of transgenes in future experiments. In addition, animals we generated in this study will be useful in experiments in which it is desirable to track cells over long periods of time in live colonies. Specifically, we expect to be able to use them to track cell migrations following the fusion of colonies with different degrees of histocompatibility. Such experiments will enable us to test previous claims regarding the cell types that migrate within chimeras, the speed and mode by which they migrate, and their ultimate fate [36,49,50].

While our study is a significant advancement for the cnidarian research community, there are several limitations that should be considered. First, we have only demonstrated successful editing at one gene, *Eef1a.* Complications may arise when targeting other loci. Second, we have made little attempt to optimize the microinjection cocktail. Our efficiency might be increased by altering the overall concentrations and/or ratios of donor template, Cas9 protein, and sgRNA. Third, we cannot rule out the possibility that the repair template has been incorporated in the *Hydractinia* genome outside *Eef1a.* In fact, given the demonstrated ease with which exogenous DNA incorporates at random into the *Hydractinia* genome, we suspect other integrations did occur but were not detectable because our transgene was designed to express a functional protein only when integrated into an existing gene in the proper reading frame. Random integrations, if they were to disrupt an endogenous gene, could alter the phenotype of a transgenic colony. Researchers using CRISPR/Cas9 genome editing to insert transgenes in the future should therefore confirm any link between an observed phenotype and the mutant genotype either by creating knockins in different genetic backgrounds, breeding to confirm that phenotypes co-segregate with the transgene, and/or probing the genomes of transgenic colonies via Southern blots or whole genome resequencing. A fourth limitation is that we observed an obvious deleterious effect in colonies expressing *Eef1a^SF-P2A-tdTom^*, but we did not determine whether this was due to toxicity of the tdTomato protein, the fusion of the SF-TAP tag to Eef1a, the SF-TAP tag itself, the potential deletion of the region downstream of *Eef1a*, or some combination of these factors. Testing these possibilities in future work will be straightforward. Finally, we were unable to immunoprecipitate Eef1a-FLAG in these studies. It is likely that insertion of a 3X FLAG tag in the *Eef1a* locus (instead of the 1X tag used in this study) would have allowed for anti-FLAG immunoprecipitation.

Beyond this work, the ability to use CRISPR/Cas9 genome editing to create targeted knockins should enable the *Hydractinia* community to study other genes of interest. It also suggests a more efficient way to screen for gene knockouts because one could design a gene trap with a fluorescent reporter, thus enabling visual screening for mutants, and germline transmitted mutations, rather than laboriously genotyping by PCR. In addition, it should now be possible to begin developing more sophisticated functional genetic tools *in Hydractinia*, including the use of systems such as Cre/lox to create inducible or conditional expression or deletion systems.

For decades, *Hydractinia* has been studied by a small but dedicated community of scientists asking fundamental biological questions. Are *Hydractinia* stem cells truly pluripotent? Are there limits to the regenerative potential of this species? How does the division of labor and polyp polymorphism evolve in colonial species? What controls colony form during ontogeny? What allows a *Hydractinia* colony to distinguish itself from other colonies so accurately? Do colonies age? We expect that the availability of high throughput genomics and genome editing will speed us toward the answers. Along the way, we hope that established cnidarian researchers and newcomers alike will join the *Hydractinia* community to ask novel questions and enrich our understanding of animal biology.

## Conclusion

This work is the first account of CRISPR/Cas9 mediated knockins in *Hydractinia* and the first example of the germline transmission of a CRISPR/Cas9 inserted transgene in a cnidarian. We expect this technology, when combined with high quality genomic resources and the logistical feasibility of genetic studies, will make *Hydractinia* a widely used model system for functional genomics.

## Materials and methods

### Animal care and breeding

*H. symbiolongicarpus* was maintained at the University of Pittsburgh. Colonies were grown on glass microscope slides and cultured in 38 liter aquaria filled with artificial seawater (Instant Ocean Reef Crystals) between 28-31 ppt and maintained at 22-23°C. Adult colonies were fed 3-day-old *Artemia* nauplii (Brine Shrimp Direct) on Monday, Wednesday, and Friday. On Tuesday and Thursday, colonies were fed a suspension of pureed oysters (fresh caught, shucked, pureed, then stored frozen). Breeding colonies were kept on a day/night cycle of 8 hrs light, 16 hrs dark. Following their first exposure to light, male and female colonies were placed in separate 3 liter bins, where they released sperm or eggs 1-1.5 hrs later. Within 20 minutes of spawning, eggs were harvested by filtering water from the female bin through a 20 µm cell strainer, and sperm was harvested by collecting 10-15 ml from the male container. Sperm and eggs were mixed together with an additional 15 ml artificial seawater in a 100 mm polystyrene petri dish, where fertilization occurred. For crosses not involving microinjection, embryos were allowed to develop into planula larvae and were settled 72-96 hours post fertilization (hpf). Metamorphosis was induced by incubating the larvae in 56 mM CsCl in filtered seawater for 4-6 hrs, and then transferring them with glass pipettes onto 25 × 75 mm glass microscope slides. Three days later, larvae that successfully metamorphosed primary polyps were hand-fed pieces of 3-day-old *Artemia* nauplii. Hand feeding continued daily until polyps were able to catch and eat whole *Artemia* nauplii on their own, whereupon they were fed identically to the adult colonies as detailed above.

### Identification and cloning of *Eef1a*

Previously published *H. symbiolongicarpus* RNA-Seq data [45] were used to identify the *Eef1a* sequence. The *Eef1a* genomic sequence was identified by BLAST searches against a draft genome of 291-10 [30]. BLAST databases were created and searched using using BLAST+ 2.4 [51]. The genomic region encoding exons 3-4 of the 295-8 *Eef1a* alleles (Additional file 2) were PCR-amplified using primers P845 and Pr846 (Additional file 1: Table S1) and Phusion High-Fidelity DNA Polymerase (ThermoFisher F530). PCR products were gel purified using the Qiagen Gel Extraction Kit (Qiagen 28706), cloned using the Blunt II TOPO Cloning Kit (Invitrogen K2800), and then sequenced.

### Genomic DNA extraction

Colonies were starved for at least two days before to DNA extraction. Tissue was harvested by scraping it from slide with a razor blade and transferring it into a 1.5 ml microcentrifuge tube, briefly centrifuging at max speed (>21,000 g), pipetting off the seawater, and storing at −80°C until extraction. To extract genomic DNA, UEB1 buffer (7M urea, 0.3125 M NaCl, 0.05 M Tris-HCl, 0.02 M EDTA, and 1% w:v N-lauroylsarcosine sodium salt) was added to the tissue, which was then ground with a pestle and incubated at 37°C for 1 hour. This was followed by a phenol/chloroform extraction and precipitation with 2.5 volumes 100% ethanol and 1/10 volume 5M sodium acetate (pH 5.2). The precipitate was pelleted, washed with 70% ethanol, and resuspended with TE. RNA was removed by adding 1/100^th^ volume of Ambion RNAse cocktail (Ambion #AM2286) and incubating at 37°C for 15 min. This was followed by another ethanol precipitation. The precipitate was pelleted, washed with 70% EtOH, resuspended with TE, and diluted to a working stock concentration of 25 ng/μl.

### *Eef1a* CRISPR target sequence selection

Before designing the repair template plasmid, a CRISPR target sequence needed to be selected. The genomic sequence of all four *Eef1a* alleles spanning primers Pr845 and Pr846 were aligned using ClustalΩ [52]. From this alignment (Additional file 3), the 3’-most PAM motif prior to the stop codon was identified and the preceding 20 nucleotides were selected as the CRISPR target sequence (sgRNA_847; Fig. 1C and Additional file 1: Table S1). To check for any obvious off-target potential, sgRNA_847 was blasted against the previously described BLAST database of the 291-10 draft genome. BLAST alignments were manually inspected for significant matches (>15 consecutive matching bases, starting at the first base prior the start of the PAM motif). Aside from the two 291-10 *Eef1a* alleles, no significant matches were found elsewhere in the genome.

### Construct design and synthesis

The Eef1a-FLAG-P2A-eGFP donor plasmid was assembled using the NEBuilder HiFi DNA Assembly kit (NEB E5520). The upstream and downstream homology arms were amplified by PCR from colony 291-10 using primers P845 and Pr846 (Additional file 1: Table S1). The *Hydractinia* codon optimized eGFP sequence was PCR amplified from pAct-I:51Act-eGFP [36]. Nucleotide sequences of the FLAG tag and P2A self cleaving peptide [53] were first codon optimized using a codon usage table generate from a publicly available *H. symbiolongicarpus* transcriptome [45] and synthesized as oligoDNA primers to be incorporated into the plasmid during Gibson assembly. The Eef1a-SF-P2A-tdTomato donor plasmid was assembled in a similar fashion. The SF-TAP sequence [46] was codon optimized and synthesized as an oligoDNA primer for assembly while a *Hydractinia* codon-optimized tdTomato sequence was amplified from a plasmid obtained from the Frank lab. These sequences were assembled and cloned using the NEBuilder HiFi DNA Assembly kit and sequenced to validate the assembly (Additional file 4). Plasmid DNA for microinjection was prepared using the GeneJET Plasmid Midiprep kit (ThermoFisher K0481), followed by an additional phenol/chloroform purification, ethanol/sodium acetate precipitation, and resuspension in 1X TE at a concentration of 1µg/µl. The plasmid was then diluted with a working buffer (20 mM HEPES; 150 mM KCl, pH 7.5) to a final concentration of 600 ng/µl, and stored at −20°C.

### Microinjection

To prepare embryos for microinjection, a cross was performed as described above. After allowing fertilization to take place in the 100 mm dish for 15 minutes, the fertilized eggs were washed three times with filtered seawater in a 50 ml conical tube to remove excess sperm and remnants of the jelly coat. Washed embryos were then loaded onto a glass microscope slide and injected until the first cell cleavage began.

The microinjection cocktail was prepared prior to injection. Recombinant Cas9 protein (PNA Bio) and a synthetic sgRNA (Synthego) were combined and incubated at 37°C for 10 minutes, after which the donor plasmid was added. The final injection cocktail consisted of 1 µg/µl Cas9, 500 ng/µl sgRNA, and 200 ng/µl donor plasmid. Immediately before injection, the cocktail was centrifuged at max speed (>21,000 g) for approx. 5 minutes to pellet precipitates and particles that would clog the injection needle. The supernatant was then loaded into the microinjection needle and injected into single-cell *Hydractinia* embryos using a FemtoJet (Eppendorf) microinjection system.

### Tissue Fixation and Immunolocalization

To prepare fixed tissue, animals were anaesthetized with menthol crystals (Sigma M2772) or 2% urethane in seawater, followed by fixation in 4% paraformaldehyde in seawater for 1 hour at room temperature (RT) or overnight (ON) at 4°C. Tissue was permeabilized by washing three times with PBS + 0.3% Triton X-100 (PBST) for 30 minutes at 4°C. For immunolocalizations, all steps were carried out at RT unless stated otherwise. Following fixation tissues were then blocked in 3% BSA/PBST (two incubations, 30 minutes each), incubated ON at 4°C with 1:1,000 dilution of mouse anti-FLAG primary antibody (Sigma F1804) in 3% BSA/PBST, washed with PBST (three washes, 10 minutes each), blocked in 3% BSA/5% goat serum/PBST for 30 min, incubated for 1 hour with a 1:1,1000 dilution of an Alexa 647 conjugated goat anti-mouse secondary antibody (ThermoFisher A-21235) in 3% BSA/5% goat serum/PBST, and washed with PBST (three washes, 10 minutes each). Nuclei were then stained ON at 4°C with 5 µg/ml DAPI, washed with PBST (three washes at 4°C, 30 minutes each), and mounted in Gelvatol mounting medium [54] prior to imaging.

### Microscopy

Images of live colonies were collected on an Olympus macroscope fitted with a 1X objective equipped with a Retiga camera and controlled by Nikon NIS Elements software. Three dimensional images of larvae collected using a Nikon A1 confocal microscope in galvanometer scanning mode with an ELWD 40x water objective (NA 1.12) with Nyquist sampling in X,Y and Z. Images were rendered and stitched using NIS elements. Confocal images of other tissues were acquired on a Nikon A1 scanning confocal microscope with 20X air objectives (NA 0.75). For multiphoton microscopy, we used a Nikon A1R MP microscope configured with the Nikon Ni-E (Nikon Instruments, Tokyo, Japan) upright motorized system. The microscope was equipped with a Chameleon Laser Vision (Coherent, Inc., CA, USA) tuned to 850nm for GFP excitation, an APO LWD 25x water immersion objective with 1.1 NA, a high-speed (30 frames/s) resonant scanning mode and a galvano scanning mode suitable for linescanning acquisitions. The Nano-Drive system was used to acquire fine high speed control of z-plane selection. The fluorescence detection unit of the microscope consisted of four detectors (photo multiplying tubes). We used detector 2 for eGFP 525/50nm (green channel). The microscope components including the laser, stage, resonant scanning head, detectors and acquisition were controlled using NIS Elements software (Nikon). Post-processing was carried out in NIS Elements software (v. 4.5) and exported as TIFF files. All adjustments were applied equally to the entire image.

### Colony area estimation

Colony area estimated by visualizing live colonies with a Leica M80 stereoscope at 10X magnification and acquiring color images in JPEG format with a Leica IC80HD camera. Images were imported into ImageJ as implemented in Fiji [55]. This was converted into a binary black and white image by adjusting the threshold level until all pixels within the colony perimeter were black and the background remained white. The “Analyzed Particles” command was then used to compute the area of the colony. The scale was determined by imaging a ruler at the same magnification. Statistical analysis was performed in GraphPad Prism.

### Genotyping

Genomic inserts were amplified using the Q5 High-Fidelity 2X Master Mix (NEB catalog M0492). To amplify each junction, PCR primers were designed so that one of the primers is tethered to the endogenous gDNA sequence of the EF1-α locus and the paired primer to the insert (Figs. 3,7). For a 50 µL PCR, 100 ng of gDNA was used. PCRs were analyzed by electrophoresis on 1% agarose gels. PCR products were purified using the GeneJET PCR Purification Kit (ThermoFisher K0701), cloned using the Blunt II TOPO Cloning Kit (Invitrogen K2800), and then sequenced.

### Protein extraction

Total protein was extracted from whole colonies of *H. symbiolongicarpus.* Prior to tissue collection, a lysis buffer consisting of RIPA buffer (Sigma R0278), 1 mM AEBSF, 1 µg/ml pepstatin A, 5 mM EDTA pH 8, 1 mM EGTA pH 8, and 1X cOmplete™ EDTA-free protease inhibitor cocktail tabs (Roche #11836170001) was prepared. Colonies were excised from glass slides, collected in eppendorf tubes, and briefly centrifuged at max speed (>21,000 g) to allow excess seawater to be removed. Then the lysis buffer was added and the tissue ground with a pestle and incubated on ice for 10-20 minutes. Lysates were then centrifuged at max speed (>21,000 g) for 10 min at 4°C. Lysate was decanted into a new tube and placed on ice. Protein was quantified using the Pierce BCA Protein Assay Kit (ThermoFisher 23225), diluted with 4x SDS-PAGE loading buffer (with either 2-mercaptoethanol or DTT), boiled for 5 minutes at 100°C, aliquoted, and stored at −80°C.

### Immunoblotting

For western blotting, a total of 10 µg of protein was run on Novex 10% Tris-Glycine mini gels (ThermoFisher XP00105BOX). Protein was transferred to a PVDF membrane (Bio-Rad 1620177) using the Pierce G2 Fast Blotter. Blots were probed according to the supplier’s recommended protocol for the following antibodies: anti-actin (Sigma A2066), anti-FLAG (Sigma F1804), and anti-StrepII (MBL International catalog #M211-3). Secondary antibodies (anti-mouse IgG-peroxidase and anti-rabbit IgG-peroxidase; Sigma A0168 and A0545, respectively) were used at 1:10,000 dilution. Signal was detected using SuperSignal West Pico Chemiluminescent Substrate kit (ThermoFisher 34078), exposed to Carestream Biomax XAR Film (Sigma 1651454), and developed using a Konica SRX-101A tabletop processor.

### Pull-down assays

Immunoprecipitations (IPs) on FLAG-P2A-eGFP expressing colonies were performed using Anti-FLAG M2 Magnetic Beads (Sigma M8823) following the supplier’s recommended protocol. For lysates from SF-P2A-tdTomato expressing colonies, pull-downs were performed using MagStrep “type3” XT magnetic beads (IBA 2-4090-002). Total protein was extracted as described above. Total volume of the lysate was approx. 1 ml, this was added to 1 µl of equilibrated beads and incubated ON at 4°C turning end-over-end. Lysate was removed and beads were washed three times at RT using 1 ml of Strep-Tactin XT Wash Buffer (IBA 2-1003-100). After final wash, beads were moved into a clean eppendorf tube and eluted by incubating beads with 30 µl Strep-Tactin XT Elution Buffer (IBA 2-1042-025) for 10 minutes at RT with periodic vortexing. Eluted protein was diluted with 4x SDS-PAGE loading buffer (with DTT), boiled for 5 minutes at 100°C, and stored at −80°C.

IPs and pull-downs were analyzed by both coomassie staining and immunoblotting (as described above). For coomassie staining, 35 µl of the eluate was run on a Novex 10% Tris-Glycine mini gel. The gel was then fixed for 10 minutes (10% acetic acid, 25% IPA) at RT, stained ON at RT in a 0.3% solution of G250 Brilliant Blue coomassie (BioRad 161-0406), destained with 10% acetic acid for approx. two hours, and then imaged.

## Abbreviations

*Eef1a*: Eukaryotic elongation factor 1 alpha
sgRNA: single guide RNA
SF-TAP: StrepII-FLAG tandem affinity purification
PCR: polymerase chain reaction; IPs - immunoprecipitations
ON: overnight
RT: room temperature
UTR: untranslated region

## Declarations

### Acknowledgements

We thank Leo Buss from Yale University for providing inbred and field-collected colonies, Sebastien Gingras from the University of Pittsburgh Transgenic and Gene Targeting Core for helpful discussions and advice, and Simon Watkins of the Center for Biological Imaging at the University of Pittsburgh for assistance with imaging. This research was supported in part by the Intramural Research Program of the National Human Genome Research Institute, National Institutes of Health.

### Availability of data and materials

Genomic contigs containing 291-10 *Eef1a* alleles have been submitted to Genbank and are available from the authors upon request. The draft *Hydractinia* genome assembly will be freely available through the *Hydractinia* Genome Project Portal, at *https://research.nhgri.nih.gov/hydractinia/.* All other data generated and analysed during this study are included in this publication (including supplemental information).

### Author Contributions

SMS and MLN conceptualized and designed the experiments. SMS performed all experiments listed in the manuscript. ZM, JMH, and BMR were responsible for animal care and maintenance. HF duplicated results in UF’s lab. SMS, MLN, GAG, and AMW performed imaging. CES and ADB provided *Hydractinia* draft genome assemblies. MLN and SMS wrote the manuscript. All authors read and approved the final manuscript.

### Ethics approval and consent to participate

Not applicable.

## Additional files

**Additional file 1. Figures S1-S3 and Table S1.**

**Additional file 2. Alignment of cloned *Eef1a* sequences from female colony 295-8.**

Two PCR amplicons (sized 2.4 kb and 2.0 kb) from the genomic region encoding the 3’ end of *Eef1a* in colony 295-8 were cloned, sequenced (3 clones each), and aligned. This revealed two alleles, with the major differences being a 52 bp and a 313 bp indel. Differences are shown in red boxes. Sequence features are shown on and above the top sequence of the alignment. Green, *Eef1a* mRNA as determined by alignments to 291-10 transcripts. Yellow, *Eef1a* coding sequence.

**Additional file 3. Alignment of *Eef1a* alleles from colonies 291-10 and 295-8.**

Nucleotide alignment of the region encoding exons 3 and 4 of *Eef1a* from the four alleles in 291-10 and 295-8. Differences are highlighted in red boxes. Sequence features are shown on and above the top sequence of the alignment. Green, mRNA as determined by alignments to 291-10 transcripts. Yellow, coding sequence. Blue, sgRNA_847 binding site.

**Additional file 4. Genbank formatted sequence of repair template pUP613 (Eef1a-FLAG-P2A-eGFP).**

**Additional file 5. Alignment of upstream and downstream integration sites from *Eef1a^FLAG-P2A-eGFP^* colonies.**

Nucleotide alignment of the regions encoding the up- and downstream integration sites from the founder colony, 347-10, and two transgenic offspring, 354-3 and 354-5. Three cloned amplicons were sequenced for each animal at each site. Primer binding sites are indicated in green on the expected sequence. Differences are highlighted in red boxes. Note that the regions that do not overlap between the upstream and downstream sequences are shown as gaps at the beginning and end of the alignment.

**Additional file 6. Z-stack of *Eef1a^FLAG-P2A-eGFP^* gastrozooid.**

Only DAPI (blue) and eGFP (green) channels are shown.

**Additional file 7. Z-stack of *Eef1a^FLAG-P2A-eGFP^* gastrozooid stained with anti-FLAG antibodies**

Same polyp as in Additional file 6. Signal from Alexa 647 (white) is shown.

**Additional file 8. Z-stack of non-transgenic gastrozooid.**

Only DAPI (blue) and eGFP (green) channels are shown.

**Additional file 9. Z-stack of non-transgenic gastrozooid stained with anti-FLAG antibodies**

Same polyp as in Additional file 8. Signal from Alexa 647 (white) is shown.

**Additional file 10. Z-stack of tentacle from *Eef1a^FLAG-P2A-eGFP^* colony.**

**Additional file 11. Z-stack of gastrozooid from *Eef1a^FLAG-P2A-eGFP^* colony.**

**Additional file 12. Z-stack of stolonal mat and base of gastrozooid from *Eef1a^FLAG-P2A-eGFP^* colony.**

**Additional file 13. Z-stack of edge of stolonal mat from *Eef1a^FLAG-P2A-eGFP^* colony.**

**Additional file 14. Z-stack of stolon from *Eef1a^FLAG-P2A-eGFP^* colony.**

**Additional file 15. Z-stack of female gonozooid head from *Eef1a^FLAG-P2A-eGFP^* colony.**

**Additional file 16. Z-stack of *Eef1a^FLAG-P2A-eGFP^* larva.**

**Additional file 17. Z-stack of non-transgenic larva.**

**Additional file 18. Genbank formatted sequence of repair template pUP660 (Eef1a-SF-P2 A-tdTom)**

**Additional file 19. Alignment of upstream and downstream integration sites from *Eef1a^SF-P2A-tdTom^* colonies.**

Nucleotide alignment of the region encoding the upstream integration site from the founder colony, 350-10, and one transgenic offspring, 355-8. Three cloned amplicons were sequenced for each animal at each site. Primer binding sites are indicated in green on the expected sequence.

